# Deucravacitinib, a tyrosine kinase 2 pseudokinase inhibitor, protects human beta cells against proinflammatory insults

**DOI:** 10.1101/2022.12.27.522037

**Authors:** Reinaldo Sousa Dos Santos, Daniel Guzman Llorens, Atenea Alexandra Perez-Serna, Angel Nadal, Laura Marroqui

**Author notes:** **Corresponding authors: Reinaldo Sousa Dos Santos**, Instituto de Investigación, Desarrollo e Innovación en Biotecnología Sanitaria de Elche, Universidad Miguel Hernández de Elche, Ed. Torregaitán, Av. de la Universidad s/n 03202 Elche, Spain., **Laura Marroqui**, Instituto de Investigación, Desarrollo e Innovación en Biotecnología Sanitaria de Elche, Universidad Miguel Hernández de Elche, Ed. Torregaitán, Av. de la Universidad s/n 03202 Elche, Spain.

## Abstract

**Aims/hypothesis:** Type 1 diabetes is characterised by pancreatic islet inflammation and autoimmune-driven pancreatic beta cell destruction. Type I interferons, such as IFNα, are key players in early human type 1 diabetes pathogenesis, as the activation of the tyrosine kinase 2 (TYK2)-signal transducer and activator of transcription (STAT) pathway induces inflammation, a long-lasting MHC class I overexpression, endoplasmic reticulum (ER) stress, and beta cell apoptosis (in synergy with IL-1β). As TYK2 inhibition has been suggested as a potential therapeutic target for the prevention or treatment of type 1 diabetes, we investigated whether the selective TYK2 inhibitor deucravacitinib could protect beta cells against the damaging effects of IFNα and other proinflammatory cytokines (i.e. IFNγ and IL-1β).

**Methods:** Inflammation, ER stress, and apoptosis were evaluated by real-time PCR, immunoblot, immunofluorescence, and nuclear dyes. The promoter activity was assessed by luciferase assay and insulin secretion and content by ELISA. All experiments were performed in the human EndoC- βH1 cell line.

**Results:** Pre-treatment with deucravacitinib prevented IFNα effects, such as STAT1 and STAT2 phosphorylation and protein expression as well as MHC class I hyperexpression, in a dose-dependent manner without affecting beta cell survival and function. Comparison between deucravacitinib and two Janus kinase inhibitors, ruxolitinib and baricitinib, showed that deucravacitinib blocked IFNα- but not IFNγ-induced signalling pathway. Pre-treatment with deucravacitinib protected beta cells from the pro-apoptotic and proinflammatory effects of two different combinations of cytokines: IFNα + IL-1β and IFNγ + IL-1β. Moreover, this TYK2 inhibitor could partially revert apoptosis and inflammation in cells previously treated with IFNα + IL-1β or IFNγ + IL-1β.

**Conclusions/interpretation:** Our findings suggest that, by protecting beta cells against the deleterious effects of proinflammatory cytokines without affecting beta cell function and survival, deucravacitinib could be repurposed for the prevention or treatment of early type 1 diabetes.

**Research in context:** What is already known about this subject?

- In type 1 diabetes, pancreatic beta cells are killed by the immune system
- In early insulitis, type I interferons are crucial for the dialogue between the immune system and pancreatic beta cells
- Activation of the TYK2-STAT pathway by IFNα induces inflammation, HLA class I overexpression, ER stress, and beta cell apoptosis.

What is the key question?

- Could the TYK2 inhibitor deucravacitinib prevent the deleterious effects of IFNα and other cytokines in beta cells?

What are the new findings?

- Deucravacitinib prevented IFNα effects in a dose-dependent manner without affecting beta cell function and survival
- Pre-treatment with deucravacitinib protected beta cells against apoptosis and inflammation induced by two different combinations of cytokines: IFNα + IL-1β and IFNγ + IL-1β
- Addition of deucravacitinib to cells pre-treated with IFNα + IL-1β or IFNγ + IL-1β partially reverted apoptosis and inflammation induced by these cytokines

How might this impact on clinical practice in the foreseeable future?

- Due to its protective effect against proinflammatory cytokines in beta cells, our findings suggest that deucravacitinib could be repurposed for the prevention or treatment of type 1 diabetes.

## Introduction

Type 1 diabetes is characterised by pancreatic islet inflammation and specific destruction of pancreatic beta cells by an autoimmune assault, which develops in the context of an inadequate “dialogue” between beta cells and the invading immune cells [1, 2].

A growing body of evidence places type I interferons (IFNs) as key players in the early stages of human type 1 diabetes pathogenesis [3]. IFNα was found in islets from type 1 diabetes patients [4–6], and laser-captured islets from living donors with recent-onset type 1 diabetes showed increased expression of IFN-stimulated genes (ISGs) [7]. In genetically susceptible children, an IFN signature was temporarily amplified preceding the development of autoantibodies and throughout the progress of type 1 diabetes [8, 9]. Recently, three type I IFN response markers, namely human MX Dynamin Like GTPase 1 (MX1), double-stranded RNA sensor protein kinase R, and HLA class I, were found to be expressed in a significantly higher percentage of insulin-containing islets from autoantibody-positive and/or recent-onset type 1 diabetes donors [10]. In human beta cells, IFNα induces inflammation, endoplasmic reticulum (ER) stress as well as a long-lasting overexpression of HLA class I via activation of the tyrosine kinase 2 (TYK2)-signal transducer and activator of transcription (STAT) pathway. Moreover, IFNα induces apoptosis in the presence of IL-1β [11–14].

Targeting the type I IFN signalling pathway has been proposed as a potential adjuvant therapy to treat at-risk individuals or patients still in the very early stages of the disease [3, 15]. Among some of the strategies that have been suggested, inhibitors of Janus kinase (JAK) proteins (JAK1-3 and TYK2) show great promise. Treatment with AZD1480 (a JAK1/JAK2 inhibitor) and ABT 317 (a JAK1-selective inhibitor) protected non-obese diabetic mice against autoimmune diabetes and reversed diabetes in newly diagnosed non-diabetic mice [16, 17]. In human beta cells, clinically used JAK inhibitors, namely ruxolitinib, cerdulatinib, and baricitinib, prevented MHC class I overexpression, ER stress, chemokine production, and apoptosis [13, 14].

Lately, attention has been focused on *TYK2*, a candidate gene for type 1 diabetes whose genetic variants that decrease TYK2 activity are associated with protection against the disease [18–20]. TYK2 is crucial for cell development and IFNα-mediated responses in human beta cells [11, 21, 22]. Partial TYK2 knockdown protected human beta cells against apoptosis and inflammation induced by polyinosinic-polycitidilic acid (poly(I:C)), a mimic of double-stranded RNA produced during viral infection [21]. In mature stem cell-islets, TYK2 knockout or pharmacological inhibition decreased T-cell-mediated cytotoxicity by preventing IFNα-induced antigen processing and presentation, including MHC class I expression [22]. As these findings place TYK2 as a critical regulator of the type I IFN signalling pathway in beta cells, selective TYK2 inhibition has emerged as a drug target to treat type 1 diabetes. Recently, two novel small molecule inhibitors binding to the TYK2 pseudokinase domain protected human beta cells against the deleterious effects of IFNα without compromising beta cell function and susceptibility to potentially diabetogenic viruses [23].

Deucravacitinib (BMS-986165), a small molecule that selectively targets the TYK2 pseudokinase domain, has shown great therapeutic potential for immune-mediated diseases, such as lupus nephritis and systemic lupus erythematosus [24, 25]. In fact, deucravacitinib has been recently approved for treatment of plaque psoriasis [26–28]. However, no preclinical studies have deeply explored the possible use of deucravacitinib in the context of type 1 diabetes. Notably, Chandra et al. recently used deucravacitinib to validate their CRISPR-Cas9-generated *TYK2* knockout in human induced pluripotent stem cells, but did not provide further characterisation of its effects in beta cells [22].

In the present study, we described the effects of deucravacitinib on the human EndoC-βH1 beta cell line, including its ability to prevent IFNα-triggered signalling pathway and subsequent damaging effects on beta cells. We report that deucravacitinib prevented IFNα effects in a dose-dependent manner without affecting beta cell survival and function. Compared with ruxolitinib and baricitinib, deucravacitinib inhibited the IFNα- but not IFNγ-stimulated signalling pathway. Interestingly, this TYK2 inhibitor protected beta cells not only against the deleterious effects of IFNα but also from other proinflammatory cytokines, namely IFNγ and IL-1β, which suggests that deucravacitinib could be introduced as an adjuvant protective therapy at different stages of the disease to avoid the progressive loss of beta cell mass.

## Methods

### Culture of EndoC-βH1 cells

The human EndoC-βH1 beta cell line [research resource identifier (RRID): CVCL_L909, Univercell-Biosolutions, France] was cultured in Matrigel/fibronectin-coated plates as previously described [29]. Cells were cultured in DMEM containing 5.6 mmol/l glucose, 10 mmol/l nicotinamide, 5.5 μg/ml transferrin, 50 μmol/l 2-mercaptoethanol, 6.7 ng/ml selenite, 2% BSA fatty acid free, 100 U/ml penicillin, and 100 μg/ml streptomycin. We confirmed that cells were mycoplasma-free using the MycoAlert Mycoplasma Detection Kit (Lonza, Basel, Switzerland).

### Cell treatments

Proinflammatory cytokine concentrations were selected according to previously established experiments in human beta cells [11, 30]: recombinant human IFNα (PeproTech Inc., Rocky Hill, NJ) at 1000 U/ml; recombinant human IFNγ (PeproTech Inc., Rocky Hill, NJ) at 1000 U/ml; and recombinant human IL-1β (R&D Systems, Abingdon, UK) at 50 U/ml. Cells were transfected with 1 μg/ml poly(I:C) (InvivoGen, San Diego, CA) as indicated [31]. Ruxolitinib, baricitinib, or deucravacitinib (Selleckchem, Planegg, Germany) were prepared by dissolution in DMSO (used as vehicle) and cells were treated as indicated in the figures. Ruxolitinib and baricitinib concentrations were selected based on previous dose-response experiments (unpublished data). For treatments involving cytokines, 2% FBS was added to the culture medium.

### Cell viability assessment

The percentage of apoptosis was measured by fluorescence microscopy upon staining with the DNA-binding dyes Hoechst 33342 and propidium iodide (Sigma-Aldrich, Saint Louis, MO, USA) as described [32]. At least 600 cells were counted for each experimental condition. Viability was assessed by two independent observers, one of whom was unaware of sample identity, with an agreement between results of >90%.

### Caspase 3/7 activity

Caspase 3/7 activity was determined using the Caspase-Glo® 3/7 assay (Promega, Madison, WI, USA) following the manufacturer’s instructions. Briefly, upon treatment in 100 μl culture medium, cells were incubated with 100 μl Caspase-Glo® 3/7 reagent at room temperature for 1 h before recording luminescence with a POLASTAR plate reader (BMG Labtech, Ortenberg, Germany).

### C-X-C motif chemokine ligand 10 (CXCL10) measurements

The release of CXCL10 to the culture medium was detected using Human ProcartaPlex immunoassays (Invitrogen, Vienna, Austria) following the manufacturer’s recommendations. Reactions were read with a MagPix system (Luminex, Austin, TX, USA).

### Luciferase reporter assays

EndoC-βH1 cells were transfected using Lipofectamine 2000 (Invitrogen, Carlsbad, CA, USA) with pRL-CMV encoding *Renilla* luciferase (Promega) and luciferase reporter constructs for either gamma-interferon activation site (GAS) (Panomics, Fremont, CA, USA) or IFN-stimulated regulatory element (ISRE) (kindly provided by Dr Izortze Santin, University of the Basque Country, Spain). After recovery, cells were treated with either IFNα for 2 h or IFNγ for 24 h [33]. Luciferase activity was measured in a POLASTAR plate reader (BMG Labtech) using the Dual-Luciferase Reporter Assay System (Promega) and corrected for the luciferase activity of the internal control plasmid, i.e. pRL-CMV.

### RNA analysis

Poly(A)^+^ mRNA was extracted using Dynabeads mRNA DIRECT kit (Invitrogen) and cDNA synthesis was performed using the High-Capacity cDNA Reverse Transcription Kit (Applied Biosystems). Real-time PCR was performed on the CFX96 Real Time System (Bio-Rad) as described [34] and the housekeeping gene ß-actin was used to correct expression values. All primers used here are listed in ESM Table 1.

### Immunoblotting and immunofluorescence analyses

Western blotting analysis was performed as described [21]. Briefly, cells were washed with cold PBS and lysed in Laemmli buffer. Immunoblot was performed using antibodies against STAT1 and STAT2 (phosphorylated and total forms; all at 1:1000 dilution), and α-tubulin (1:5000). Peroxidase-conjugated antibodies (1:5000) were used as secondary antibodies. SuperSignal West Femto chemiluminescent substrate (Thermo Scientific, Rockford, IL, USA) and ChemiDoc XRS+ (Bio-Rad Laboratories, Hercules, CA, USA) were used to detect bands.

Immunofluorescence was carried out as described [11, 21]. First, cells were washed with cold PBS and fixed with 4% paraformaldehyde. Afterwards, cells were permeabilised and incubated with the mouse anti-MHC Class I (W6/32) antibody (1:1000). The Alexa Fluor 568 polyclonal goat anti-mouse IgG (1:500) was used as secondary antibody. Upon staining with Hoechst 33342, coverslips were mounted with fluorescent mounting medium (Dako, Carpintera, CA, USA) and immunofluorescence was observed with an inverted fluorescence microscope Zeiss Confocal

LSM900 with Airyscan 2 microscope equipped with a camera (Zeiss-Vision, Munich, Germany), and images were acquired at x40 magnification and analysed using ZEN software (version 3.3; Zeiss-Vision, Munich, Germany) and open-source FIJI software (version 2.0; https://fiji.sc). All antibodies used here are provided in ESM Table 2.

### Glucose-stimulated insulin secretion

After preincubation in modified Krebs-Ringer for 1 h, cells were sequentially stimulated with low (0 mmol/l) and high glucose (20 mmol/l) for 1 h (each stimulation) as previously described [35]. Insulin secreted and insulin content from lysed cells were measured using a human insulin ELISA kit (Mercodia, Uppsala, Sweden) following the manufacturer’s instructions. See ESM Methods for further details.

### Statistical analyses

The GraphPad Prism 7.0 software (GraphPad Software, La Jolla, CA, USA) was used for statistical analyses. Data are shown as mean ± SEM of independent experiments (i.e. considering EndoC- βH1 cells from different passages as *n* = *1*, with individual data added to the column bars). The statistical significance of differences between groups was evaluated using one-way ANOVA followed by Dunnett’s test or two-way ANOVA followed by Sidak’s test or by Dunnett’s test, as appropriate.

## Results

### Deucravacitinib prevents IFNα effects in EndoC-βH1 cells

IFNα-mediated JAK/TYK2 activation leads to phosphorylation of STAT1 and STAT2, which will eventually upregulate several ISGs, including *STAT1/2, HLA-ABC, CXCL10*, and *MX1* (ESM Fig. 1a). Pre-treatment of EndoC-βH1 cells with deucravacitinib inhibited IFNα-induced STAT1 and STAT2 phosphorylation in a dose-dependent manner, where deucravacitinib showed greater potency towards IFNα-stimulated STAT1 inhibition (Fig. 1a,b). We then selected two doses, 10 and 1000 nmol/l, for the follow-up experiments. Next, we examined the effect of deucravacitinib on the kinetics of IFNα-induced STAT activation. As expected, the phosphorylation of STAT1 and STAT2 was markedly amplified by IFNα at early time points (1–4 h) and returned to baseline by 24 h (Fig. 1c,d and ESM Fig. 1b). STAT1 and STAT2 protein expression augmented in a time dependent manner. Although both proteins were already upregulated by 8 h, STAT2 reached an expression peak at 16 h, while STAT1 expression was still increasing by 24 h (Fig. 1c-f). Exposure to 1000 nmol/l deucravacitinib abrogated the IFNα-stimulated STAT1 and STAT2 phosphorylation and protein expression, whereas 10 nmol/l deucravacitinib had only a minor effect (Fig. 1c-f and ESM Fig. 1b). These findings are better evidenced by analysing the area under the curve of the phosphorylated and total forms of STAT1 and STAT2 (ESM Fig. 1c-f). Finally, MHC class I protein expression stimulated by IFNα was completely blocked by 1000 nmol/l deucravacitinib (Fig. 1g,h).

**Figure 1.**
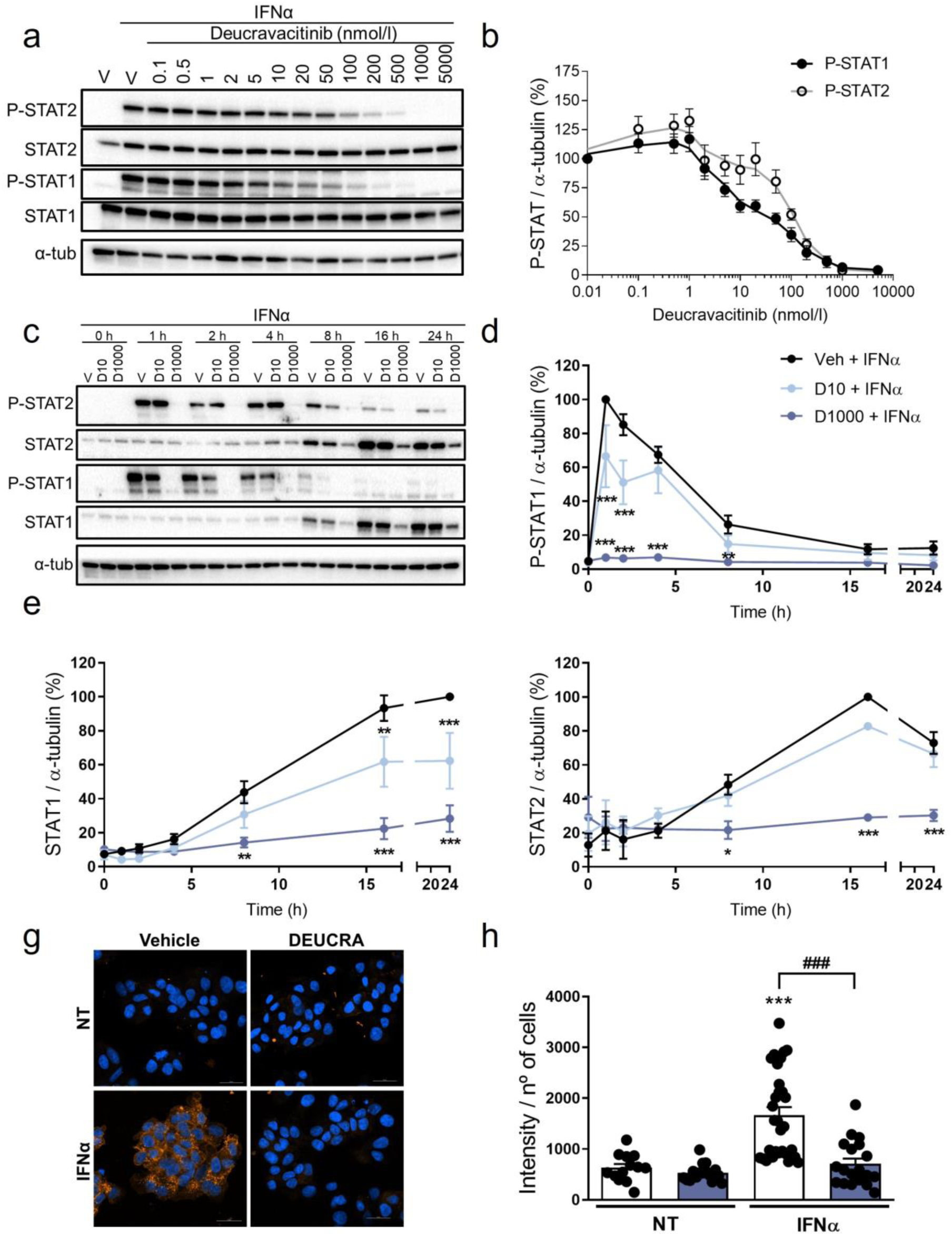
Deucravacitinib inhibits IFNα-mediated STAT phosphorylation and MHC class I overexpression. (**a**,**b**) EndoC-βH1 cells were treated with vehicle (V) or pre-treated with the indicated deucravacitinib concentrations for 1 h. Afterwards, cells were left untreated or treated with IFNα (1000 U/ml) in the absence or presence of deucravacitinib for 1 h. (**a**, **b**) Protein expression was measured by western blot. (**a**) Images are representative of four to six independent experiments. (**b**) Densitometry results are shown for P-STAT1 (black circles) and P-STAT2 (white circles). Values were normalised to α-tubulin, and then to the value of IFNα alone of each experiment (considered as 100%). (**c**-**h**) EndoC-βH1 cells were treated with vehicle (V or Veh, black circles) or pre-treated with deucravacitinib (10 [D10, soft blue circles] and 1000 nmol/l [D1000, dark blue circles]) for 1 h. Afterwards, cells were left untreated or treated with IFNα (1000 U/ml) in the absence or presence of deucravacitinib for 1–24 h (**c**-**f**) or 24 h (**g**, **h**). (**c**-**f**) Protein expression was measured by western blot. (**c**) Images are representative of three to six independent experiments. (**d**-**f**) Densitometry results are shown for P-STAT1 (**d**), STAT1 (**e**), and STAT2 (**f**). Values were normalised to α-tubulin, and then to the highest value of each experiment (considered as 1). (**g**, **h**) Immunocytochemistry analysis of MHC class I (red) and Hoechst 33342 (blue) upon exposure to IFNα in the absence (white bars) or presence of 1000 nmol/l deucravacitinib (dark blue bars) for 24 h. Representative images of three independent experiments (13–30 images/coverslip) (**g**) and quantification (**h**) are shown. Data are means ± SEM of three to six independent experiments. (**d**-**f**) \**p*≤0.05, \*\**p*≤0.01 and \*\*\**p*≤0.001 vs Vehicle + IFNα (two-way ANOVA plus Dunnett’s test). (**h**) \*\*\**p*<0.001 vs the respective untreated (NT) (two-way ANOVA plus Sidak’s test); ###*p*≤0.001, as indicated by bars (two-way ANOVA plus Dunnett’s test).

### Beta cell survival and function are not affected by deucravacitinib

We examined whether deucravacitinib would interfere with beta cell survival and function in the absence or presence of IFNα. After 24 h exposure, 1000 nmol/l deucravacitinib did not affect beta cell viability (ESM Fig. 2a) nor change glucose-stimulated insulin secretion and insulin content (ESM Fig. 2b,c). IFNα did not affect beta cell viability and function as described [11, 23].

**Figure 2.**
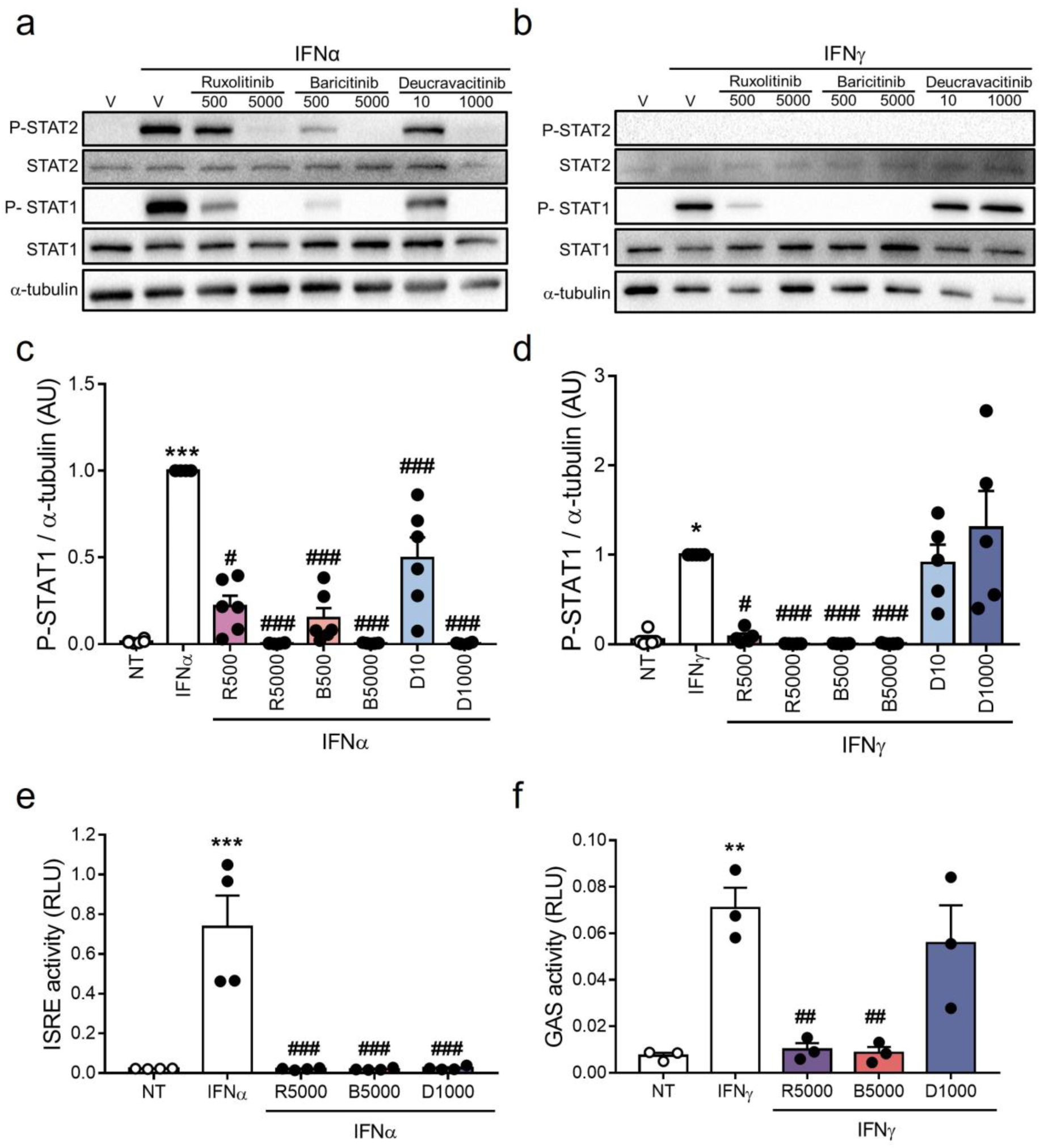
Deucravacitinib blocks IFNα- but not IFNγ-induced pathway. (**a**-**d**) EndoC-βH1 cells were treated with vehicle (V, white bars) or pre-treated with ruxolitinib (500 and 5000 nmol/l; R500 and R5000), baricitinib (500 and 5000 nmol/l; B500 and B5000), or deucravacitinib (10 and 1000 nmol/l; D10 and D1000) for 1 h. Afterwards, cells were left untreated (NT, white circles) or treated with either IFNα (1000 U/ml) (**a**, **c**) or IFNγ (1000 U/ml) (**b**, **d**) in the absence or presence of each inhibitor for 1 h. (**a**-**d**) Protein expression was measured by western blot. (**a**, **b**) Images are representative of five to six independent experiments. (**c**, **d**) Densitometry results are shown for P-STAT1. Values were normalised to α-tubulin, and then to the value of IFNα (**c**) or IFNγ alone (**d**) of each experiment (considered as 1). (**e**, **f**) EndoC-βH1 cells were transfected with a pRL-CMV plasmid (used as internal control) plus either ISRE (**e**) or GAS (**f**) promoter reporter constructs. After 48 h of recovery, cells were treated with vehicle (white bars) or pre-treated with ruxolitinib (5000 nmol/l; R5000), baricitinib (5000 nmol/l; B5000), or deucravacitinib (1000 nmol/l; D1000) for 1 h. Afterwards, cells were left untreated (NT, white circles) or treated with either IFNα (1000 U/ml) for 2 h (**e**) or IFNγ (1000 U/ml) for 24 h (**f**) in the absence or presence of each inhibitor. Relative luciferase units (RLU) were measured by a luminescent assay. Data are means ± SEM of three to six independent experiments. \**p*≤0.05, \*\**p*≤0.01 and \*\*\**p*≤0.001 vs the respective untreated (NT) (one-way ANOVA plus Dunnett’s test). #*p*≤0.05, ##*p*≤0.01 and ###*p*≤0.001 vs IFNα (**c**, **e**) or IFNγ (**d**, **f**) (one-way ANOVA plus Dunnett’s test).

### IFNα, but not IFNγ signalling pathway is blocked by deucravacitinib

We compared deucravacitinib with ruxolitinib and baricitinib, two JAK1/JAK2 inhibitors previously tested in beta cells [13, 14, 36]. First, we measured STAT1 and STAT2 phosphorylation upon stimulation with IFNα or IFNγ (Fig. 2a-d and ESM Fig. 3a,b). Ruxolitinib, baricitinib, and deucravacitinib prevented IFNα-stimulated increase in P-STAT1 and P-STAT2 levels (Fig. 2a,c and ESM Fig. 3b). Nevertheless, deucravacitinib did not change IFNγ-induced STAT1 phosphorylation, whereas ruxolitinib and baricitinib blocked it (Fig. 2b,d). Of note, IFNγ did not induce STAT2 phosphorylation (Fig. 2b) [33]. Next, we assessed ISRE and GAS activities upon stimulation with IFNα and IFNγ (Fig. 2e,f and ESM Fig. 3c,d). All three inhibitors abrogated IFNα-stimulated ISRE reporter activity (Fig. 2e), whereas IFNγ-induced GAS activation was barely affected by deucravacitinib (Fig. 2f). As expected, ISRE and GAS activities were not stimulated by, respectively, IFNγ and IFNα (ESM Fig. 3c,d). These findings corroborate the deucravacitinib mode of action, which specifically binds to and inhibits TYK2 without affecting JAK1/JAK2 pathways. As TYK2 does not participate in the IFNγ pathway, the lack of deucravacitinib effect in IFNγ-stimulated changes is expected.

**Figure 3.**
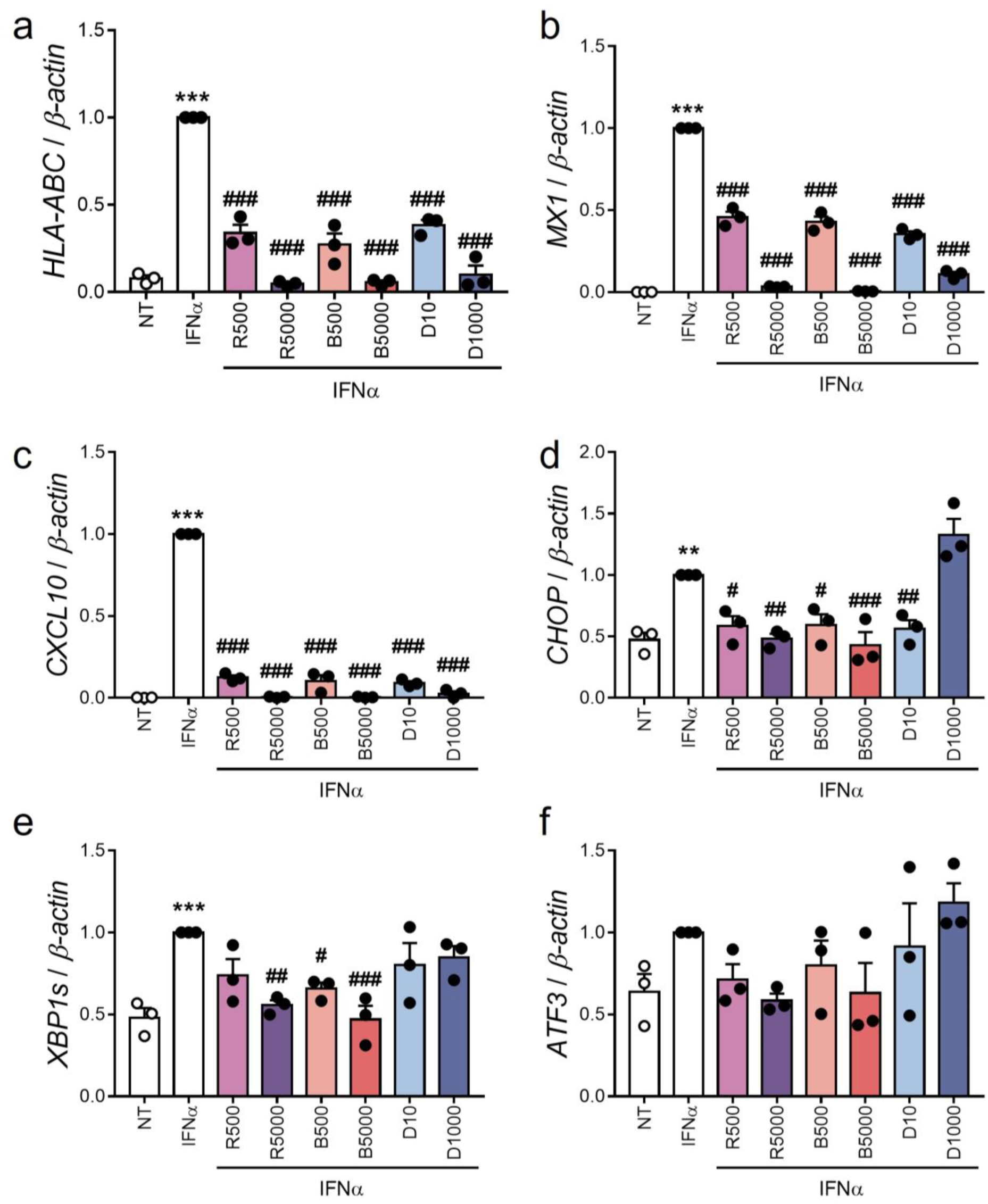
Deucravacitinib abrogates IFNα-induced expression of ISGs but not ER stress markers. (**a**-**f**) EndoC-βH1 cells were treated with vehicle (white bars) or pre-treated with ruxolitinib (500 and 5000 nmol/l; R500 and R5000), baricitinib (500 and 5000 nmol/l; B500 and B5000), or deucravacitinib (10 and 1000 nmol/l; D10 and D1000) for 1 h. Afterwards, cells were left untreated (NT) or treated with IFNα (1000 U/ml) in the absence or presence of each inhibitor for 24 h. mRNA expression of *HLA-ABC* (**a**), *MX1* (**b**), *CXCL10* (**c**), *CHOP* (**d**), *XBP1s* (**e**), and *ATF3* (**f**) was analysed by real-time PCR, normalised to β-actin and then to the value of IFNα alone of each experiment (considered as 1). Data are means ± SEM of three independent experiments. \*\**p*≤0.01 and \*\*\**p*≤0.001 vs the respective untreated (NT) (one-way ANOVA plus Dunnett’s test). #*p*≤0.05, ##*p*≤0.01 and ###*p*≤0.001 vs IFNα (one-way ANOVA plus Dunnett’s test).

### Deucravacitinib blocks IFNα-induced upregulation of ISGs, but not ER stress markers

We assessed the effects of these three inhibitors on the expression of some ISGs and ER stress markers. All three inhibitors prevented IFNα-induced upregulation of *HLA-ABC, CXCL10*, and *MX1* in a dose-dependent manner (Fig. 3a-c). Although ruxolitinib and baricitinib inhibited the expression of the ER stress markers C/EBP homologous protein (*CHOP*, also known as *DDIT3*) and spliced isoform of XBP1 X-box binding protein 1 (*XBP1s*), only the lower dose of deucravacitinib reduced CHOP expression (Fig. 3d,e). None of the three inhibitors changed the mRNA expression of activating transcription factor 3 (*ATF3*) (Fig. 3f).

### Deucravacitinib prevents IFNα + IL-1β-induced effects in beta cells

Previous studies showed that a combination of IFNα + IL-1β, two cytokines that might be present in the islet milieu at early stages of insulitis, induces beta cell apoptosis, inflammation, and ER stress [11, 14, 23] (Fig. 4a). Then, we investigated whether deucravacitinib protects beta cells after IFNα + IL-1β exposure for 24 h. We observed that deucravacitinib completely prevented IFNα + IL-1β-induced apoptosis, which was assessed by two approaches, namely DNA-binding dyes and caspase 3/7 activity assay (Fig. 4b,c). Moreover, cells treated with deucravacitinib showed reduced levels of P-STAT1 and STAT1 (Fig. 4d-f) as well as *HLA-ABC, MX1, CXCL10*, and *CHOP* mRNA expression (Fig. 4g-j). MHC class I protein expression and CXCL10 secretion to the medium were also decreased by TYK2 inhibition (Fig. 4k-m).

**Figure 4.**
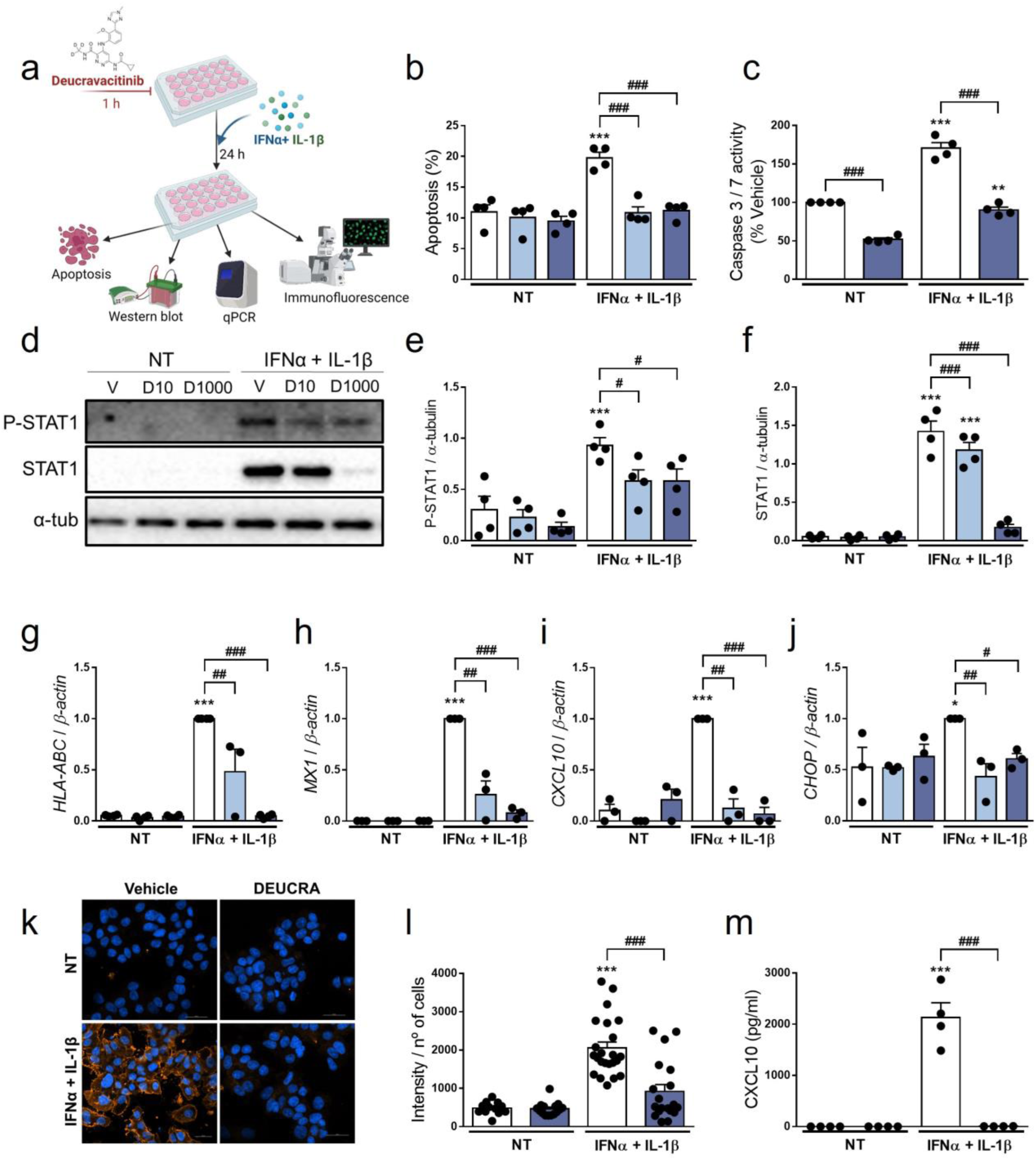
Pre-treatment with deucravacitinib prevents IFNα + IL-1β effects. (**a**) Experimental design of the pre-treatment with deucravacitinib and subsequent exposure to IFNα + IL-1β for 24 h. EndoC-βH1 cells were treated with vehicle (V, white bars) or pre-treated with deucravacitinib (10 [D10, soft blue bars] and 1000 nmol/l [D1000, dark blue bars]) for 1 h. Afterwards, cells were left untreated (NT) or treated with IFNα + IL-1β (1000 U/ml + 50 U/ml, respectively) in the absence or presence of deucravacitinib for 24 h. (**b**) Apoptosis was evaluated using Hoechst 33342/propidium iodide staining. (**c**) Caspase 3/7 activity was measured by a luminescent assay. Results are expressed as % vehicle-treated cells in the absence of cytokines (NT). (**d**-**f**) Protein expression was measured by western blot. (**d**) Images are representative of four independent experiments. Densitometry results are shown for P-STAT1 (**e**) and P-STAT2 (**f**). Values were normalised to α-tubulin. (**g**-**j**) mRNA expression of *HLA-ABC* (**g**), *CXCL10* (**h**), *MX1* (**i**), and *CHOP* (**j**) was analysed by real-time PCR, normalised to β-actin and then to the value of Vehicle treated with IFNα + IL-1β (considered as 1). (**k**, **l**) Immunocytochemistry analysis of MHC class I (red) and Hoechst 33342 (blue) upon exposure to IFNα + IL-1β in the absence (white bars) or presence of deucravacitinib (dark blue bars) for 24 h. Representative images of three independent experiments (12–23 images/coverslip) (**k**) and quantification (**l**) are shown. (**m**) CXCL10 secreted to the medium was determined by ELISA. Data are means ± SEM of three to four independent experiments. \*\**p*≤0.01 and \*\*\**p*≤0.001 vs the respective untreated (NT) (two-way ANOVA plus Sidak’s test). #*p*≤0.05, ##*p*≤0.01 and ###*p*≤0.001, as indicated by bars (two-way ANOVA plus Dunnett’s test).

### Deucravacitinib abrogates IFNγ + IL-1β-induced apoptosis in beta cells

We then evaluated whether deucravacitinib would protect against cytokines that, as compared with IFNα, probably appear later in the progression of islet inflammation, such as IFNγ and IL-1β [1, 37] (Fig. 5a). After treatment for 24 h, deucravacitinib inhibited IFNγ + IL-1β-induced apoptosis in a dose-dependent manner (60% and 92% protection at 10 and 1000 nmol/l, respectively) (Fig. 5b). These results were confirmed by the caspase 3/7 activity assay (Fig. 5c). Pre-treatment with deucravacitinib did not block IFNγ + IL-1β-induced STAT1 phosphorylation and protein expression (Fig. 5d-f) or *HLA-ABC* mRNA expression (Fig. 5g). In fact, P-STAT1 levels were higher in cells treated with 1000 nmol/l deucravacitinib than in vehicle-treated cells (Fig. 5d,e). Conversely, deucravacitinib diminished *MX1* and *CXCL10* mRNA expression, whereas *CHOP* was reduced at the lower dose of deucravacitinib (Fig. 5h-j).

**Figure 5.**
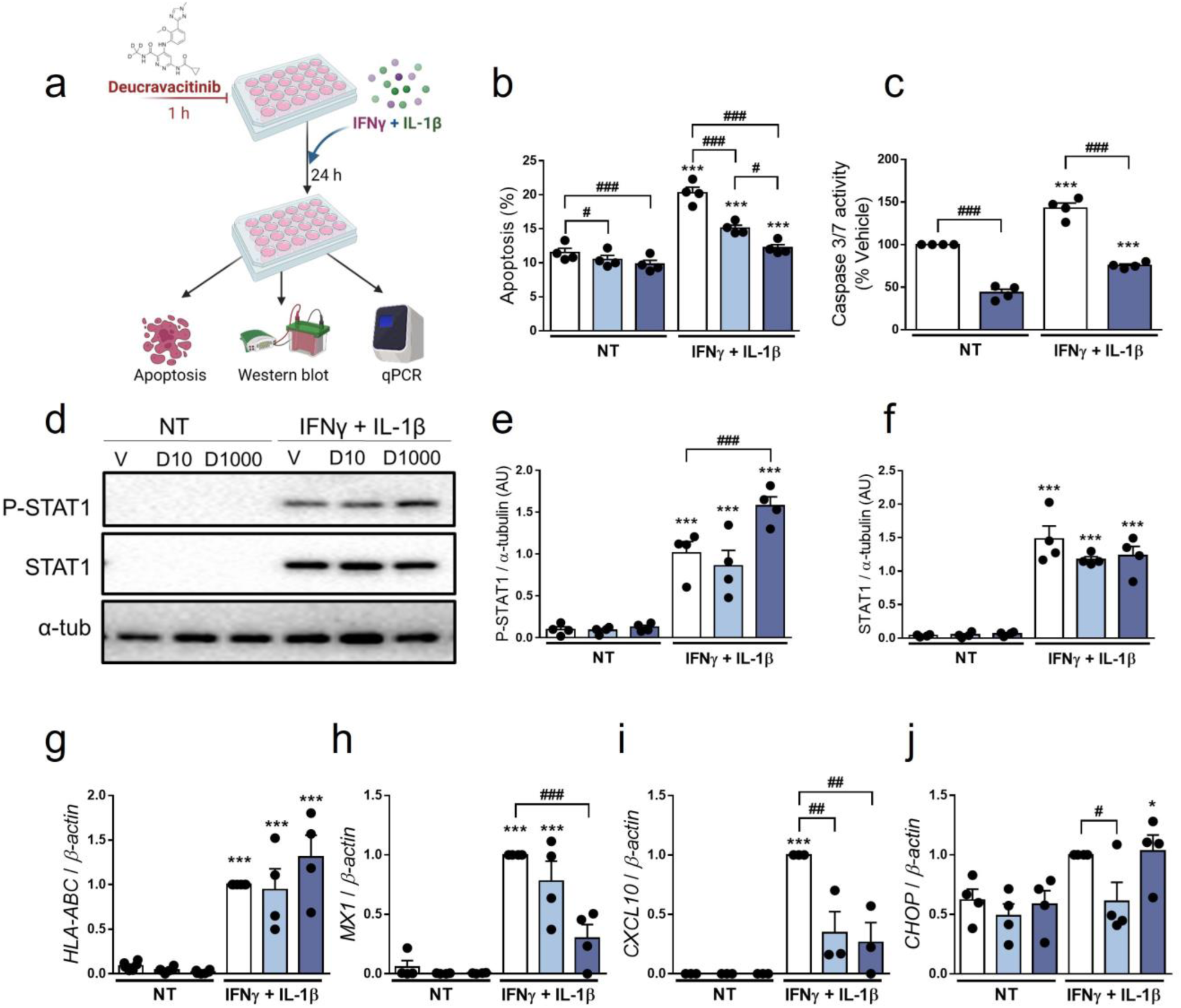
Pre-treatment with deucravacitinib prevents IFNγ + IL-1β effects. (**a**) Experimental design of the pre-treatment with deucravacitinib and subsequent exposure to IFNγ + IL-1β for 24 h. EndoC-βH1 cells were treated with vehicle (V, white bars) or pre-treated with deucravacitinib (10 [D10, soft blue bars] and 1000 nmol/l [D1000, dark blue bars]) for 1 h. Afterwards, cells were left untreated (NT) or treated with IFNγ + IL-1β (1000 U/ml + 50 U/ml, respectively) in the absence or presence of deucravacitinib for 24 h. (**b**) Apoptosis was evaluated using Hoechst 33342/propidium iodide staining. (**c**) Caspase 3/7 activity was measured by a luminescent assay. Results are expressed as % vehicle-treated cells in the absence of cytokines (NT). (**d**-**f**) Protein expression was measured by western blot. Images are representative of four independent experiments (**d**). Densitometry results are shown for P-STAT1 (**e**) and P-STAT2 (**f**). Values were normalised to α-tubulin. (**g**-**j**) mRNA expression of *HLA-ABC* (**g**), *CXCL10* (**h**), *MX1* (**i**), and *CHOP* (**j**)was analysed by real-time PCR, normalised to β-actin and then to the value of Vehicle treated with IFNγ + IL-1β (considered as 1). Data are means ± SEM of three to four independent experiments. \**p*≤0.05, \*\**p*≤0.01 and \*\*\**p*≤0.001 vs the respective untreated (NT) (two-way ANOVA plus Sidak’s test). #*p*≤0.05, ##*p*≤0.01 and ###*p*≤0.001, as indicated by bars (two-way ANOVA plus Dunnett’s test).

### Poly(I:C)-induced apoptosis is not changed by TYK2 inhibition

As intracellular exposure to poly(I:C), a mimic of viral infection, results in the production and secretion of type I IFNs as well as beta cell apoptosis [21, 38, 39], we tested deucravacitinib after treatment with poly(I:C) for 24 h. Contrary to what we observed for cytokine-triggered cell death (Figs 4 and 5), apoptosis induced by poly(I:C) was not altered by deucravacitinib (ESM Fig. 4a). Curiously, despite the absence of an effect on viability, deucravacitinib inhibited poly(I:C)- stimulated *HLA-ABC, MX1*, and *CHOP* upregulation (ESM Fig. 4b-d). While *CXCL10* mRNA expression was dampened by TYK2 inhibition, CXCL10 protein secretion was not significantly reduced (ESM Fig. 4e,f).

### The harmful effects of cytokines are partially reverted by deucravacitinib

Up to this point, we investigated whether pre-treatment with deucravacitinib would prevent the effects of different cytokines in beta cells. Here, we assessed if deucravacitinib could reverse these damaging effects. EndoC-βH1 cells were pre-treated with either IFNα + IL-1β or IFNγ + IL-1β for 24 h. Afterwards, 1000 nmol/l deucravacitinib was added for an additional 24 h still in the presence of the different mixes of cytokines (Figs 6a and 7a). IFNα + IL-1β-induced apoptosis was partially reversed by deucravacitinib (60% decrease) (Fig. 6b). The expression of *HLA-ABC* mRNA stimulated by IFNα + IL-1β remained unchanged in the presence of deucravacitinib (Fig. 6c), which agrees with previous data showing a long-lasting expression of *HLA-ABC* [13]. STAT1 protein levels and CXCL10 secretion as well as *CHOP* mRNA expression were reduced by 26%- 42%, while the expression of *MX1* and *CXCL10* was completely inhibited by deucravacitinib (Fig. 6 d-i). Of note, STAT1 phosphorylation was not detected, probably due to a process of desensitisation following the initial IFNα stimulus [33] (Fig. 6c).

**Figure 6.**
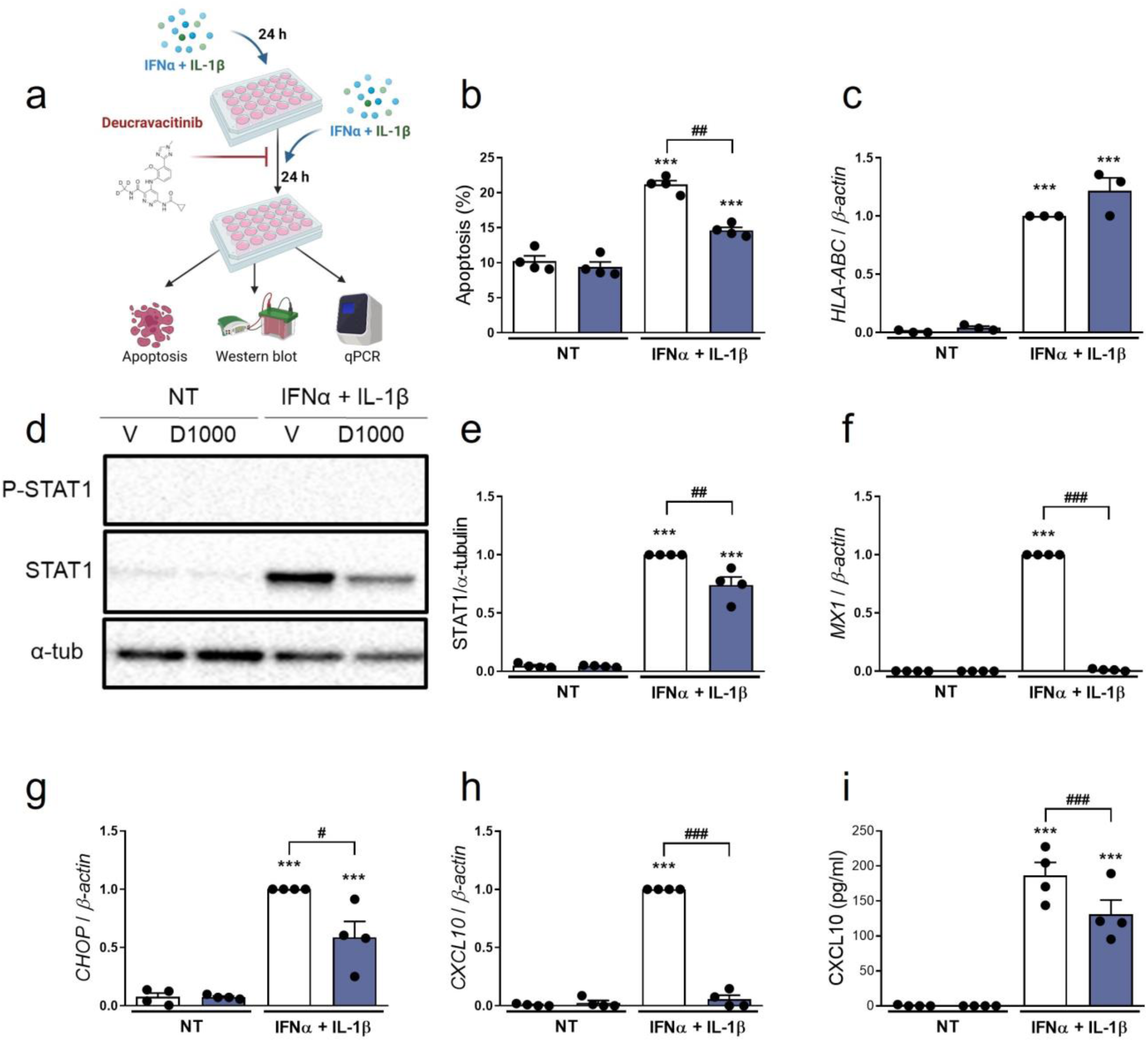
Treatment with deucravacitinib partially reverts IFNα + IL-1β-induced changes. (**a**) Experimental design of the pre-treatment with IFNα + IL-1β and subsequent exposure to IFNα + IL-1β in the presence of deucravacitinib for 24 h. EndoC-βH1 cells were left untreated (NT) or pre-treated with IFNα + IL-1β (1000 U/ml + 50 U/ml, respectively) for 24 h. Afterwards, cells were treated with vehicle (V, white bars) or 1000 nmol/l deucravacitinib (D1000, dark blue bars) in the absence (NT) or presence of IFNα + IL-1β for 24 h. (**b**) Apoptosis was evaluated using Hoechst 33342/propidium iodide staining. (**c**, **f**-**h**) mRNA expression of *HLA-ABC* (**c**), *MX1* (**f**), *CHOP* (**g**), and *CXCL10* (**h**) was analysed by real-time PCR, normalised to β-actin and then to the value of Vehicle treated with IFNα + IL-1β (considered as 1). (**d**, **e**) Protein expression was measured by western blot. (**d**) Images are representative of four independent experiments. (**e**) Densitometry results are shown for STAT1. Values were normalised to α-tubulin. (**i**) CXCL10 secreted to the medium was determined by ELISA. Data are means ± SEM of three to four independent experiments. \*\**p*≤0.01 and \*\*\**p*≤0.001 vs the respective untreated (NT) (two-way ANOVA plus Sidak’s test). #*p*≤0.05, ##*p*≤0.01 and ###*p*≤0.001, as indicated by bars (two-way ANOVA plus Dunnett’s test).

**Figure 7.**
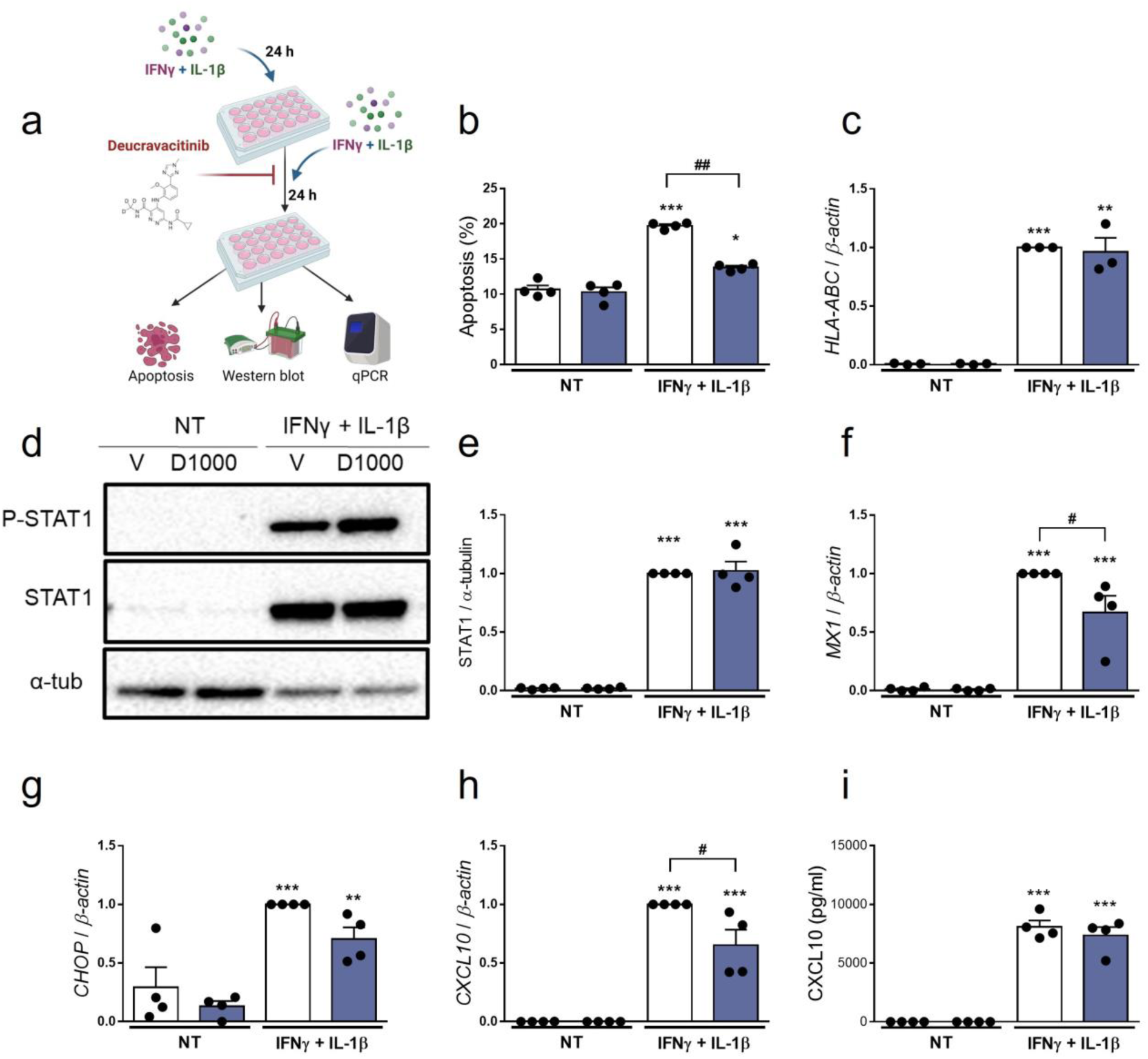
Treatment with deucravacitinib partially reverts IFNγ + IL-1β-induced changes. (**a**) Experimental design of the pre-treatment with IFNγ + IL-1β and subsequent exposure to IFNγ + IL-1β in the presence of deucravacitinib for 24 h. EndoC-βH1 cells were left untreated (NT) or pre-treated with IFNγ + IL-1β (1000 U/ml + 50 U/ml, respectively) for 24 h. Afterwards, cells were treated with vehicle (V, white bars) or 1000 nmol/l deucravacitinib (D1000, dark blue bars) in the absence (NT) or presence of IFNγ + IL-1 β for 24 h. (**b**) Apoptosis was evaluated using Hoechst 33342/propidium iodide staining. (**c**, **f**-**h**) mRNA expression of *HLA-ABC* (**c**), *MX1* (**f**), *CHOP* (**g**), and *CXCL10* (**h**) was analysed by real-time PCR, normalised to β-actin and then to the value of Vehicle treated with IFNγ + IL-1β (considered as 1). (**d**, **e**) Protein expression was measured by western blot. (**d**) Images are representative of four independent experiments. (**e**) Densitometry results are shown for STAT1. Values were normalised to α-tubulin. (**i**) CXCL10 secreted to the medium was determined by ELISA. Data are means ± SEM of three to four independent experiments. \**p*≤0.05, \*\**p*≤0.01 and \*\*\**p*≤0.001 vs the respective untreated (NT) (two-way ANOVA plus Sidak’s test). #*p*≤0.05 and ##*p*≤0.01, as indicated by bars (two-way ANOVA plus Dunnett’s test).

Similarly to IFNα + IL-1β (Fig. 6), deucravacitinib diminished IFNγ + IL-1β-induced apoptosis (64% decrease) but did not modify *HLA-ABC* mRNA expression (Fig. 7b,c). Protein levels of STAT1 and CXCL10, however, were not altered by the TYK2 inhibitor, whereas a slight, non-significant 30% reduction was seen in *CHOP* expression (Fig. 7d,e,g,i). Expression of *MX1* and *CXCL10* was only partially blocked by deucravacitinib under IFNγ + IL-1β conditions (Fig. 7f,h).

## Discussion

Due to its implication in type 1 diabetes pathogenesis, IFNα pathway has arisen as an interesting therapeutic target. Reduction of IFNα extracellular levels, blockade of IFNα itself and/or its receptor, and reduction of the activity of proteins mediating IFN effects have been proposed as means to diminish IFNα deleterious effects [40]. In rodent models, some of these strategies have successfully prevented diabetes development [16, 17, 41–43].

One of the most promising therapeutic approaches for type 1 diabetes prevention/early treatment is the targeting of the JAK-STAT pathway with JAK inhibitors [3, 15]. This strategy has been clinically approved for the treatment of some autoimmune diseases, including systemic lupus erythematosus [44], rheumatoid arthritis [45], and psoriasis [26, 27]. Although there are no approved JAK inhibitors being clinically used for type 1 diabetes, recent preclinical data suggest that these inhibitors could be repurposed to treat this disease [13, 14, 16, 17, 22, 23]. In fact, a clinical trial aiming to determine whether baricitinib could slow the progressive, immune-mediated loss of beta cell function and mass that occurs in type 1 diabetes is currently ongoing [46].

In the current study, we tested whether the TYK2 inhibitor deucravacitinib could protect human beta cells against the deleterious effects of IFNα and other cytokines. We chose to focus on this TYK2 inhibitor for two main reasons: first, due to the importance of TYK2 to type 1 diabetes pathogenesis. For instance, TYK2 regulates pro-apoptotic and proinflammatory pathways via regulation of the IFNα signalling, antigen processing and presentation, and modulation of cytokine and chemokine production in beta cells [21, 22]. Second, exploring a drug that has been recently approved by the US Food and Drug Administration to treat another autoimmune disease, namely plaque psoriasis [28], increases its repositioning potential for type 1 diabetes and facilitates the bench-to-bedside transition.

Deucravacitinib is a small-molecule ligand that binds to and stabilizes the TYK2 pseudokinase domain, leading to highly potent and selective allosteric TYK2 inhibition [24, 47]. Inhibition of IFNα-induced STAT phosphorylation by deucravacitinib has been shown in several cell types, such as CD3^+^ T cells, CD19^+^ B cells, and CD14^+^ monocytes [24]. Here we showed that deucravacitinib also prevents IFNα-stimulated STAT1 and STAT2 phosphorylation in human EndoC-βH1 cell line. Furthermore, in agreement with previous findings [24], deucravacitinib also showed higher potency against TYK2-mediated phosphorylation of STAT1 compared with STAT2 phosphorylation in our experimental model. Notably, at the concentrations used in our study, deucravacitinib did not affect beta cell function and viability, which is a desired feature for a drug with therapeutic potential.

Compared with ruxolitinib and baricitinib, two clinically available JAK1/JAK2 inhibitors, deucravacitinib was more potent against IFNα-stimulated STAT phosphorylation, ISRE activity, and mRNA expression of *HLA-ABC, MX1*, and *CXCL10*. However, unlike ruxolitinib and baricitinib, deucravacitinib did not block the IFNα-mediated upregulation of the ER stress markers *CHOP* and *XBP1s*. Our results partially agree with the ones reported by Coomans de Brachène et al., where two TYK2 inhibitors failed to prevent IFNα-induced *CHOP* expression in EndoC-βH1 cells but inhibited *CHOP* and *ATF3* expression in dispersed human islets [23].

Prior studies have shown that other JAK/TYK2 inhibitors could prevent the detrimental effects of IFNα + IL-1β, such as apoptosis and inflammation [14, 23]. Therefore, we proceeded to investigate whether deucravacitinib could protect beta cells against the harmful effects of two different combinations of cytokines: IFNα + IL-1β (early insulitis) and IFNγ + IL-1β (late insulitis). In both scenarios, pre-treatment with deucravacitinib not only protected against cytokine-induced apoptosis but prevented the upregulation of *HLA-ABC*, *MX1*, *CXCL10*, and *CHOP*. Additionally, in cells treated with IFNα + IL-1β, deucravacitinib blocked the overexpression of MHC class I at the cell surface and CXCL10 secretion to the medium. Importantly, the addition of deucravacitinib when cytokine exposure was already ongoing could reverse, at least in part, the deleterious effects of these cytokines. Although it seems clear that deucravacitinib confers protection against IFNα + IL-1β by directly inhibiting the TYK2-mediated pathway, it remains to be answered how deucravacitinib protects against IFNγ + IL-1β-induced effects. Indeed, our present data suggest that deucravacitinib does not interfere with the IFNγ- mediated signalling pathway.

Due to TYK2 role in the IFNα-mediated antiviral response, as evidenced by the regulation of *MX1* mRNA expression and STAT1 protein levels [11, 22, 23], another desired characteristic for a TYK2 inhibitor would be to block IFNα pathway without sensitising beta cells to viral infections. To test this hypothesis, we mimicked a viral infection by exposing EndoC-βH1 cells to poly(I:C). In our model, deucravacitinib abrogated poly(I:C)-stimulated inflammation but did not change poly(I:C)-induced beta cell apoptosis. Even though our results were obtained with a synthetic viral double-stranded RNA analogue, they conform to a previous study showing that two novel TYK2 inhibitors did not sensitize human beta cells to potentially diabetogenic coxsackieviruses CVB1 and CVB5 [23]. Importantly, deucravacitinib and these two TYK2 inhibitors share the same mechanism of action, i.e. binding to the TYK2 pseudokinase domain.

Based on our findings, it will be interesting to test whether novel small molecule TYK2 pseudokinase ligands [48] could also protect beta cells from IFNα deleterious effects. Nevertheless, we must bear in mind that completely inhibiting TYK2 may be counterproductive, as it might lead to susceptibility to microorganisms (e.g. mycobacteria and virus) and immunodeficiency [49]. Thus, regardless of the TYK2 inhibitor chosen, we should focus on doses that induce a partial inhibition, as seen in individuals with a protective single nucleotide polymorphism in the *TYK2* gene [18], as it could offer maximal efficacy with reduced risk of developing secondary infections.

One potential limitation of our study is its purely in vitro nature, which may limit our conclusions regarding the use of deucravacitinib to treat a disease as complex as type 1 diabetes. Conversely, our findings, along with those reported by Coomans de Brachène et al. [23] and Chandra et al. [22], provide further preclinical evidence that TYK2 inhibitors could be considered a strategy for an early therapy for type 1 diabetes. The next logical step would be to investigate whether our in vitro findings could be translated to animal models of type 1 diabetes (e.g. NOD and RIP-B7.1 mice).

In conclusion, we have provided evidence that deucravacitinib protects beta cells against the deleterious effects of proinflammatory cytokines, such as IFNα, IFNγ and IL-1β, without affecting beta cell function and survival. Our present findings add to the growing body of evidence showing that TYK2 inhibition may be an efficient strategy to treat type 1 diabetes. Moreover, we suggest that deucravacitinib could be repurposed to treat pre-symptomatic type 1 diabetes subjects (i.e. positive for 2–3 autoantibodies but still normoglycemic) or be introduced in the early stages of type 1 diabetes onset.

## Abbreviations

ATF3: Activating transcription factor 3
CHOP: C/EBP homologous protein
CXCL10: C-X-C motif chemokine ligand 10
ER: Endoplasmic reticulum
GAS: Gamma-interferon activation site
ISG: IFN-stimulated genes
ISRE: IFN-stimulated regulatory element
JAK: Janus kinase
MX1: MX Dynamin Like GTPase 1
PKR: Double-stranded RNA sensor protein kinase R
Poly(I:C): Polyinosinic-polycytidylic acid
STAT: Signal transducer and activator of transcription
TYK2: Tyrosine kinase 2
XBP1: X-box binding protein 1
XBP1s: Spliced isoform of XBP1

## Acknowledgements

The authors are grateful to Beatriz Bonmati Botella, Maria Luisa Navarro, and Salomé Ramon for their excellent technical support. Once again, we thank Dr. Izortze Santin, University of the Basque Country, Spain, for providing the luciferase reporter construct for ISRE. Figures 4a, 5a, 6a, 7a, ESM Fig. 1a, and ESM Fig. 3a were created with BioRender.com.

## Data availability

All data generated or analysed during this study are included in this published article (and its supplementary information files). Data are however available from the corresponding authors upon reasonable request.

## Authors’ relationships and activities

The authors declare that there are no relationships or activities that might bias, or be perceived to bias, their work.

## Contribution statement

RSDS and LM: Conceptualization, Supervision, Visualization, Investigation, Formal analysis, and Writing – original draft & editing; DGL and AAP-S: Investigation, Formal analysis, and Writing – review & editing; AN: Resources and Writing – review & editing; LM: Resources, Funding acquisition, and Project administration. All authors have read and given approval to the final version of the manuscript. RSDS and LM are guarantors of this work and, as such, had full access to all the data presented herein and take responsibility for the integrity of the data and the accuracy of data analyses.

