## Supplementary Material for "Deucravacitinib, a tyrosine kinase 2 pseudokinase inhibitor, protects human beta cells against proinflammatory insults"

**Electronic Supplementary Material**

### ESM Methods

#### Glucose-stimulated insulin secretion

*Insulin secretion*: EndoC-βH1 cells (70,000 cells/well) were incubated in modified Krebs-Ringer buffer (115 mmol/l NaCl, 24 mmol/l $NaHCO_3$, 5 mmol/l KCl, 1 mmol/l $CaCl_2$, 1 mmol/l $MgCl_2$, 10 mmol/l HEPES pH 7.4, and 0.1% BSA) for 1 h before glucose stimulation. Upon starvation, cells were sequentially incubated with low (0 mmol/l) and high glucose (20 mmol/l) for 1 h (each incubation). After each stimulatory period, the incubation medium was collected, placed onto ice, and centrifuged for 5 min at 700 *g* (4°C). The supernatant was transferred into a new, fresh tube and stored at -20°C until insulin measurements. The amount of secreted insulin as *% of total insulin* was calculated as previously described [1]. Data are normalised to insulin secretion at 20 mmol/l glucose in vehicle-treated cells without IFNα (considered as 100%).

*Insulin content*: upon incubation with low and high glucose, cells were lysed in a cell lysis solution containing 137 mmol/l NaCl, 0.1% Triton X100, 1% glycerol, 2 mmol/l EGTA, 20 mmol/l Tris pH 8.0, and protease inhibitor cocktail. Cell lysates were centrifuged for 5 min at 700 *g* (4°C); the supernatant was transferred into a new, fresh tube and stored at -20°C until insulin measurements. Insulin content (ng insulin/$10^6$ cells) was normalised to the condition Vehicle untreated (NT) (considered as 100%).

Insulin secreted and insulin content were measured using a human insulin ELISA kit (Mercodia, Uppsala, Sweden).

| | **Forward** | **Reverse** |
|---|---|---|
| | **Sequence (5'-3')** | **Sequence (5'-3')** |
| ***β-actin*** | CTGTACGCCAACACAGTGCT | GCTCAGGAGGAGCAATGATC |
| ***HLA-ABC*** | GAGAACGGGAAGGAGACGC | CATCTCAGGGTGAGGGGCT |
| ***CXCL10*** | GTGGCATTCAAGGAGTACCTC | GCCTTCGATTCTGGATTCAG |
| ***MX1*** | AGACAGGACCATCGGAATCT | GTAACCCTTCTTCAGGTGGAAC |
| ***ATF3*** | GCTGTCACCACGTGCAGTAT | TTTGTGTTAACGCTGGGAGA |
| ***CHOP*** | AACGGAAACAGAGTGGTCATT | GCTTGAGCCGTTCATTCTCT |
| ***XBP1s*** | CCGCAGCAGGTGCAGG | GAGTCAATACCGCCAGAATCCA |

**ESM Table 1. Primers used in the present study.**

| Antibody | Manufacturer | Catalogue number | Species raised in | Dilution | RRID |
|---|---|---|---|---|---|
| Phospho-STAT1 | Cell Signaling Technology | 9167 | Rabbit, monoclonal | 1:1000 | AB_561284 |
| Phospho-STAT2 | Cell Signaling Technology | 88410 | Rabbit, monoclonal | 1:1000 | AB_2800123 |
| STAT1 | Cell Signaling Technology | 9172 | Rabbit, polyclonal | 1:1000 | AB_2198300 |
| STAT2 | Santa Cruz Biotechnology | sc-514193 | Mouse, monoclonal | 1:1000 | AB_2810271 |
| MHC Class I (W6/32) | Enzo Life Sciences | ALX-805-711-C100 | Mouse, monoclonal | 1:1000 | AB_11179235 |
| α-Tubulin | Sigma | T9026 | Mouse, monoclonal | 1:5000 | AB_477593 |
| Goat anti-mouse IgG | Bio-rad | 170-6516 | Goat, polyclonal | 1:5000 | AB_11125547 |
| Goat anti-rabbit IgG | Bio-rad | 170-6515 | Goat, polyclonal | 1:5000 | AB_11125142 |
| Alexa Fluor 568 goat anti-mouse IgG | Invitrogen | A-11031 | Goat, polyclonal | 1:500 | AB_144696 |

**ESM Table 2. Antibodies used in the present study.**

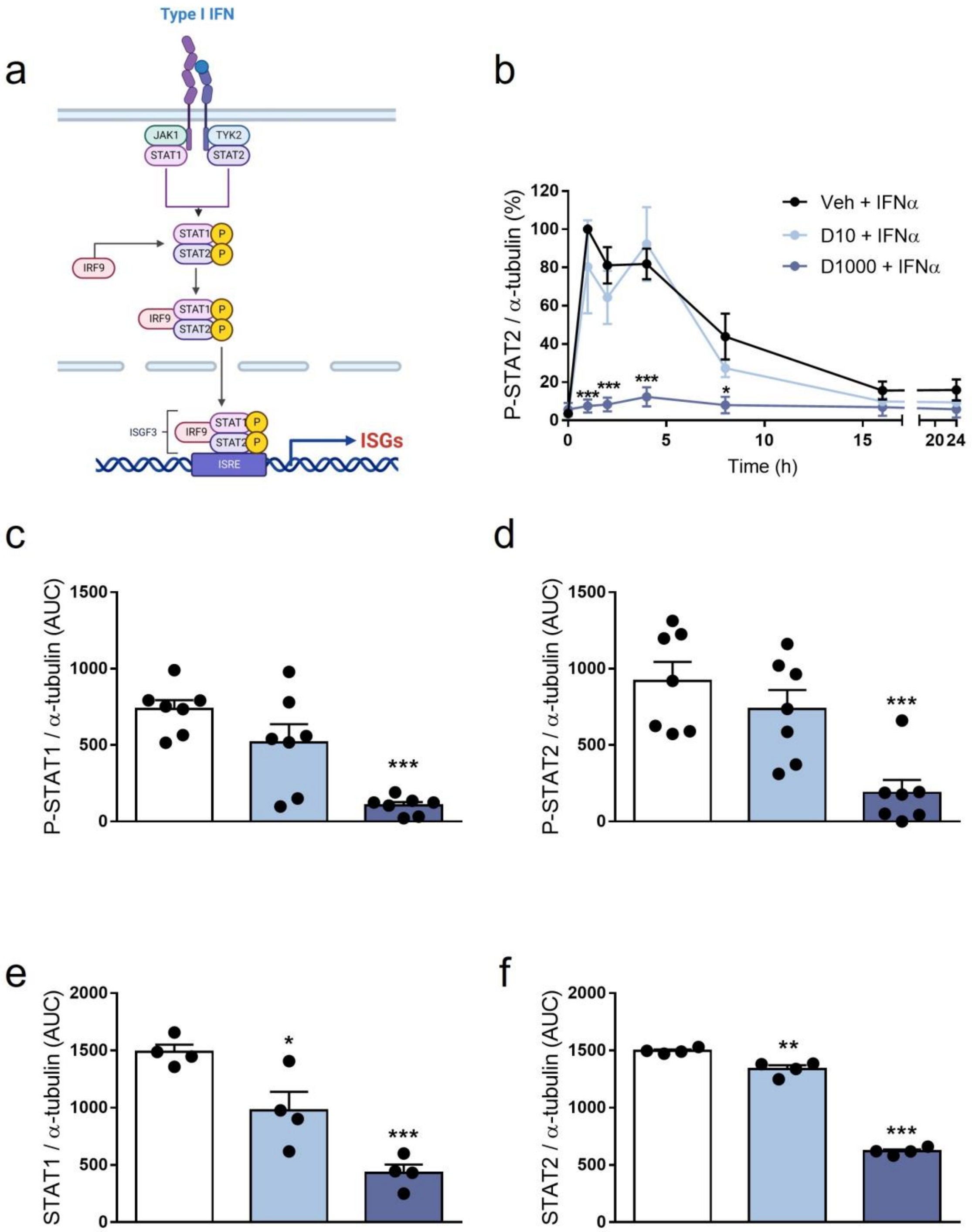


**ESM Figure 1. Deucravacitinib prevents IFNα-induced STAT1/2 phosphorylation and expression.** (**a**) Schematic representation of the type I interferon pathway. JAK1, Janus kinase 1; TYK2, Tyrosine kinase; IFNAR, IFNα/β receptor. EndoC-βH1 cells were treated with vehicle (Veh, black circles or white bars) or pre-treated with deucravacitinib (10 [D10, soft blue circles or bars] and 1000 nmol/l [D1000, dark blue circles or bars]) for 1 h. Afterwards, cells were left untreated (NT) or treated with IFNα (1000 U/ml) in the absence or presence of deucravacitinib for 1-24 h. (**b-f**) Protein expression was measured by western blot. Images representative of three to six independent experiments are shown in Fig. 1c. (**b**) Densitometry results are shown for P-STAT2. Values were normalised to α-tubulin, and then to the highest value of each experiment (considered as 1). (**c**-**f**) Quantification of the area under curve (AUC) of P-STAT1 (**c**), P-STAT1 (**d**), STAT1 (**e**), and STAT2 (**f**). Data are means ± SEM of three to four independent experiments. (**b**) ***$p \leq 0.001$ vs Vehicle + IFNα (two-way ANOVA plus Dunnett's test). (**c**-**f**) *$p<0.05$ and ***$p<0.001$ vs Vehicle + IFNα (one-way ANOVA plus Dunnett's test).

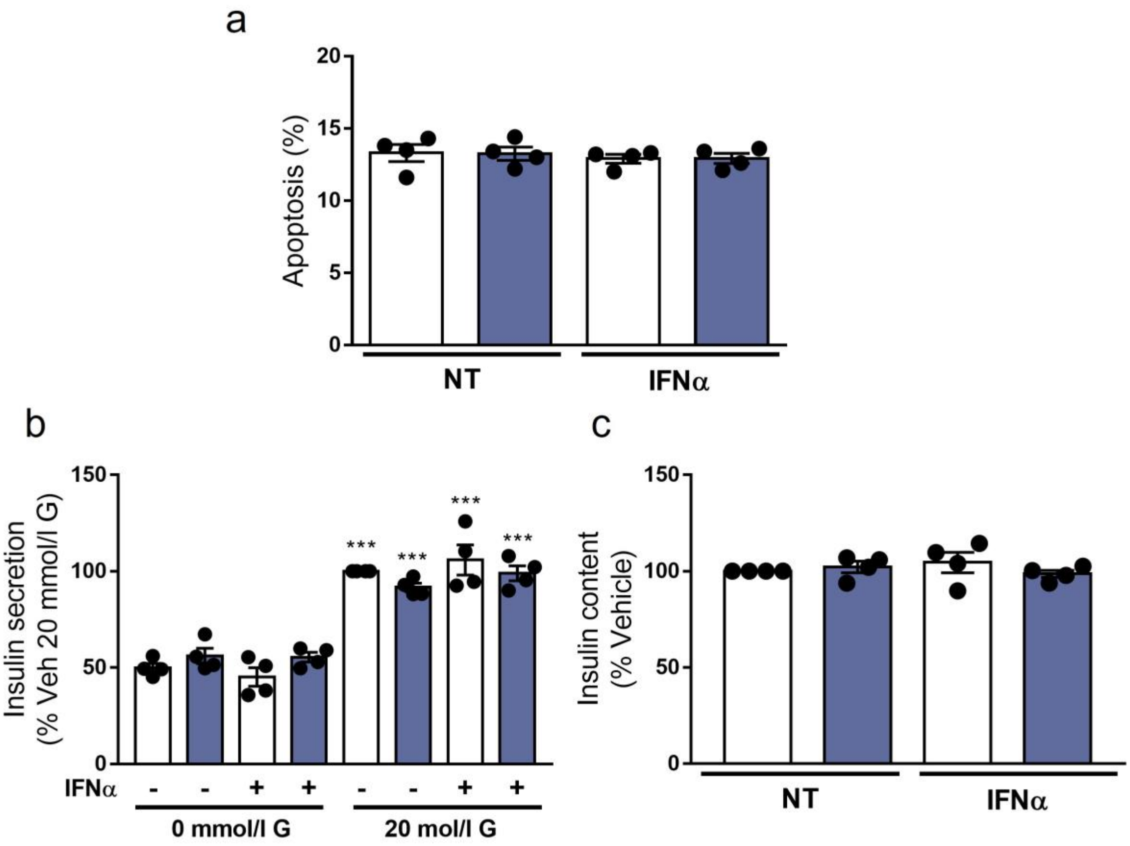


**ESM Figure 2. Deucravacitinib does not change beta cell function and survival.** EndoC-βH1 cells were treated with vehicle (white bars) or pre-treated with 1000 nmol/l deucravacitinib (dark blue bars) for 1 h. Afterwards, cells were left untreated (NT) or treated with IFNα (1000 U/ml) in the absence or presence of deucravacitinib for 24 h. (**b**) Apoptosis was evaluated using Hoechst 33342/propidium iodide staining. (**b, c**) Glucose-stimulated insulin secretion (**b**) and insulin content (**c**) were measured by ELISA. Data are means ± SEM of four independent experiments. ***$p \leq 0.001$ vs the respective treatment at 0 mmol/l glucose (G) (two-way ANOVA plus Sidak's test).

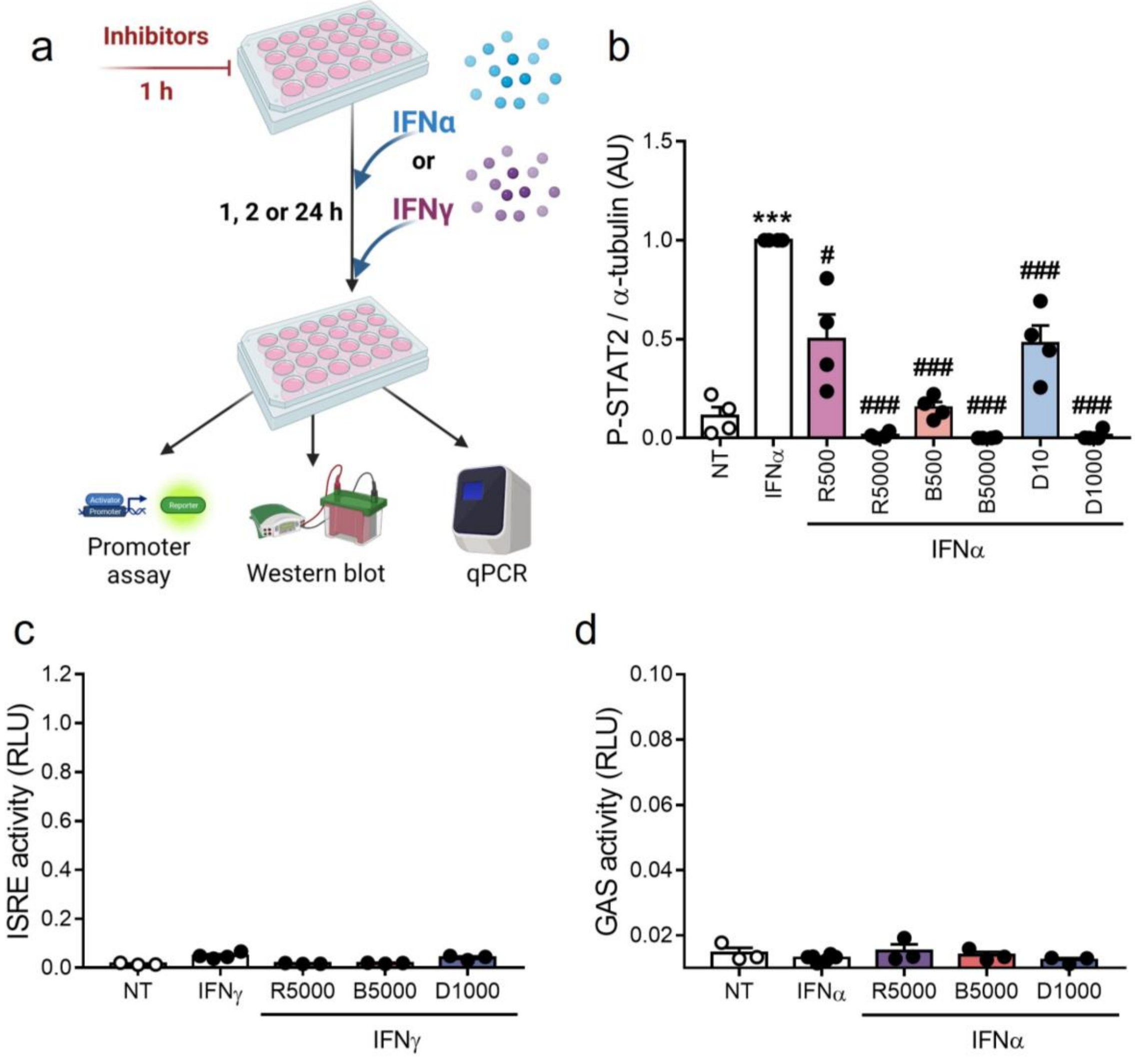


**ESM Figure 3. JAK/TYK2 inhibitors prevent STAT2 phosphorylation.** (**a**) Experimental design of the pre-treatment with deucravacitinib and subsequent exposure to IFNα or IFNγ for 1, 2 or 24 h. EndoC-βH1 cells were treated with vehicle (white bars) or pre-treated with ruxolitinib (500 and 5000 nmol/l; R500 and R5000), baricitinib (500 and 5000 nmol/l; B500 and B5000), or deucravacitinib (10 and 1000 nmol/l; D10 and D1000) for 1 h. Afterwards, cells were left untreated (NT, white circles) or treated with either IFNα (1000 U/ml) or IFNγ (1000 U/ml) in the absence or presence of each inhibitor for 1 h. (**b**) Protein expression was measured by western blot. Images representative of five to six independent experiments are shown in Fig. 2a. Densitometry results are shown for P-STAT2. Values were normalised to α-tubulin, and then to the value of IFNα of each experiment (considered as 1). (**c**, **d**) EndoC-βH1 cells were transfected with a pRL-CMV plasmid (used as internal control) plus either ISRE (**c**) or GAS (**d**) promoter reporter constructs. After 48 h of recovery, cells were treated with vehicle (white bars) or pre-treated with ruxolitinib (5000 nmol/l; R5000), baricitinib (5000 nmol/l; B5000), or deucravacitinib (1000 nmol/l; D1000) for 1 h. Afterwards, cells were left untreated (NT, white circles) or treated with either IFNγ (1000 U/ml) for 2 h (**c**) or IFNα (1000 U/ml) for 24 h (**d**) in the absence or presence of each inhibitor. Relative luciferase units (RLU) were measured by a luminescent assay. Data are means ± SEM of three to four independent experiments. $^{***}p \leq 0.001$ vs untreated (NT) (one-way ANOVA plus Dunnett's test). $^{\#}p \leq 0.05$ and $^{\#\#\#}p \leq 0.001$ vs IFNα (one-way ANOVA plus Dunnett's test).

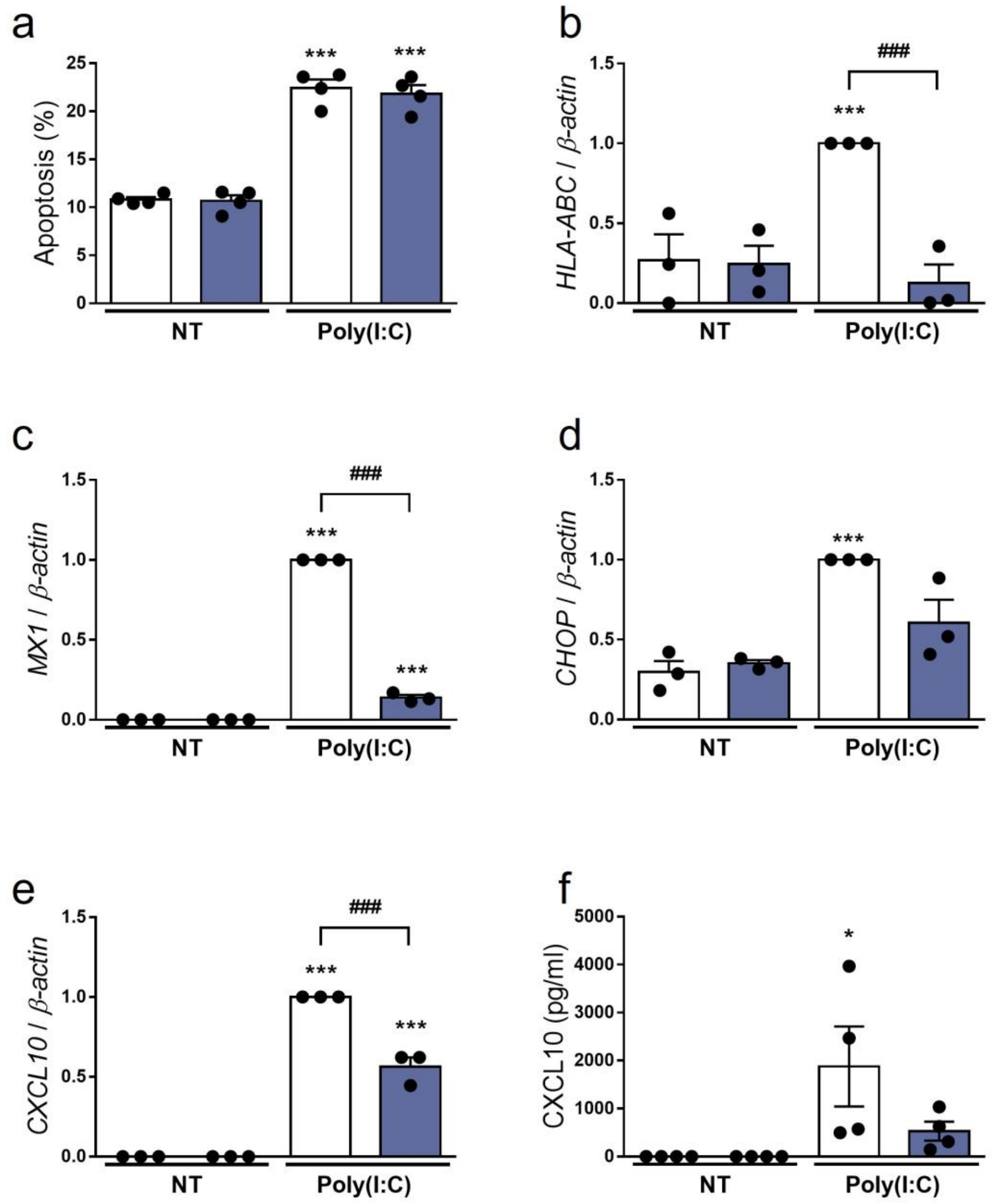


**ESM Figure 4. Deucravacitinib prevents poly(I:C)-induced inflammation but not apoptosis.** EndoC-βH1 cells were treated with vehicle (white bars) or pre-treated with 1000 nmol/l deucravacitinib (dark blue bars) for 1 h. Afterwards, cells were left untreated (NT) or treated with intracellular poly(I:C) (1 µg/ml) in the absence or presence of deucravacitinib for 24 h. (**a**) Apoptosis was evaluated using Hoechst 33342/propidium iodide staining. (**b**-**e**) mRNA expression of *HLA-ABC* (**b**), *MX1* (**c**), *CHOP* (**d**), and *CXCL10* (**e**) was analysed by real-time PCR, normalised to β-actin and then to the value of Vehicle treated with poly(I:C) (considered as 1). (**f**) CXCL10 secreted to the medium was determined by ELISA. Data are means ± SEM of four independent experiments. $^{**}p\leq0.01$ and $^{***}p\leq0.001$ vs the respective untreated (NT) (two-way ANOVA plus Sidak's test). $^{\#\#}p\leq0.01$ and $^{\#\#\#}p\leq0.001$, as indicated by bars (two-way ANOVA plus Dunnett's test).
